# Large-scale estimation of bacterial and archaeal DNA prevalence in metagenomes reveals biome-specific patterns

**DOI:** 10.1101/2024.05.16.594470

**Authors:** Raphael Eisenhofer, Antton Alberdi, Ben J. Woodcroft

**Author notes:** these authors contributed equally (Raphael Eisenhofer, Ben J. Woodcroft).

## Abstract

Metagenomes often contain many reads derived from eukaryotes. However, there is usually no reliable method for estimating the prevalence of non-microbial reads in a metagenome, forcing many analysis techniques to make the often-faulty assumption that all reads are microbial. For instance, the success of metagenome-assembled genome (MAG) recovery efforts is assessed by the number of reads mapped to recovered MAGs, a procedure which will underestimate the true fidelity if eukaryotic reads are present. Here we present “SingleM microbial_fraction” (SMF), a scalable algorithm that robustly estimates the number of bacterial and archaeal reads in a metagenome, and the average microbial genome size. SMF does not use eukaryotic reference genome data and can be applied to any Illumina metagenome. Based on SMF, we propose the “Domain-Adjusted Mapping Rate” (DAMR) as an improved metric to assess microbial genome recovery from metagenomes. We benchmark SMF on simulated and real data, and demonstrate how DAMRs can guide genome recovery. Applying SMF to 136,284 publicly available metagenomes, we report substantial variation in microbial fractions and biome-specific patterns of microbial abundance, providing insights into how microorganisms and eukaryotes are distributed across Earth. Finally, we show that substantial amounts of human host DNA sequence data have been deposited in public metagenome repositories, possibly counter to ethical directives that mandate screening of these reads prior to release. As the adoption of metagenomic sequencing continues to grow, we foresee SMF being a valuable tool for the appraisal of genome recovery efforts, and the recovery of global patterns of microorganism distribution.

## Introduction

Shotgun sequencing is a common technique used to study DNA extracted from microbial habitats of all kinds, both host-associated and environmental (1). In many habitats, microbial organisms (here Bacteria and Archaea) are numerically dominant, and give rise to the majority of reads sequenced. However, for many samples, it is not clear at the outset of the experiment how much of the DNA is microbial. DNA might be derived from other elements of the community such as fungi, protozoa, or viruses. In host-associated microbiomes, DNA might also be derived from a host’s genome, diet, symbionts or parasites (2). Metagenomic analysis techniques typically assume that the fraction of reads derived from non-microbial sources is negligible. For instance, read-centric techniques ascribe functions to all reads, and tools for estimating taxonomic profiles from metagenomic data may use reference databases composed only of Bacteria and Archaea (3). This simplifying assumption is usually made as a consequence of our inability to accurately assess the quantity and identity of non-microbial reads in metagenomic datasets. How often this assumption holds is an open question.

Genome-resolved metagenomics is a strategy that is increasingly employed to study microbiomes (4, 5). In this approach, DNA sequences (typically short reads) are assembled into larger contiguous sequences (contigs), which are then sorted (binned) using nucleotide frequencies and/or read coverage information into metagenome-assembled genomes (MAGs) (6–8). This powerful approach can generate near-complete genomes from diverse sample types — providing genomic information for uncultured microbial diversity (9). Despite significant progress in bioinformatic methodology, however, recovered genomes rarely represent the entirety of the sequenced community.

The standard metric for evaluating genome recovery efforts is to map metagenomic reads against the recovered MAGs (10). If the vast majority of reads map, then genome recovery has been successful. In contrast, a low proportion of reads mapping can indicate that the microbial community is not well represented by the MAGs. This poor representation might be due to insufficient coverage for assembly or other challenges e.g. high population heterogeneity interfering with assembly. However, the presence of non-microbial reads in a metagenome will also reduce read mapping rates. If non-microbial reads are sufficiently numerous, genome recovery efforts will be falsely judged as poor by the standard metric even if they represent the entire microbial community (**Figure 1**).

**Figure 1:**
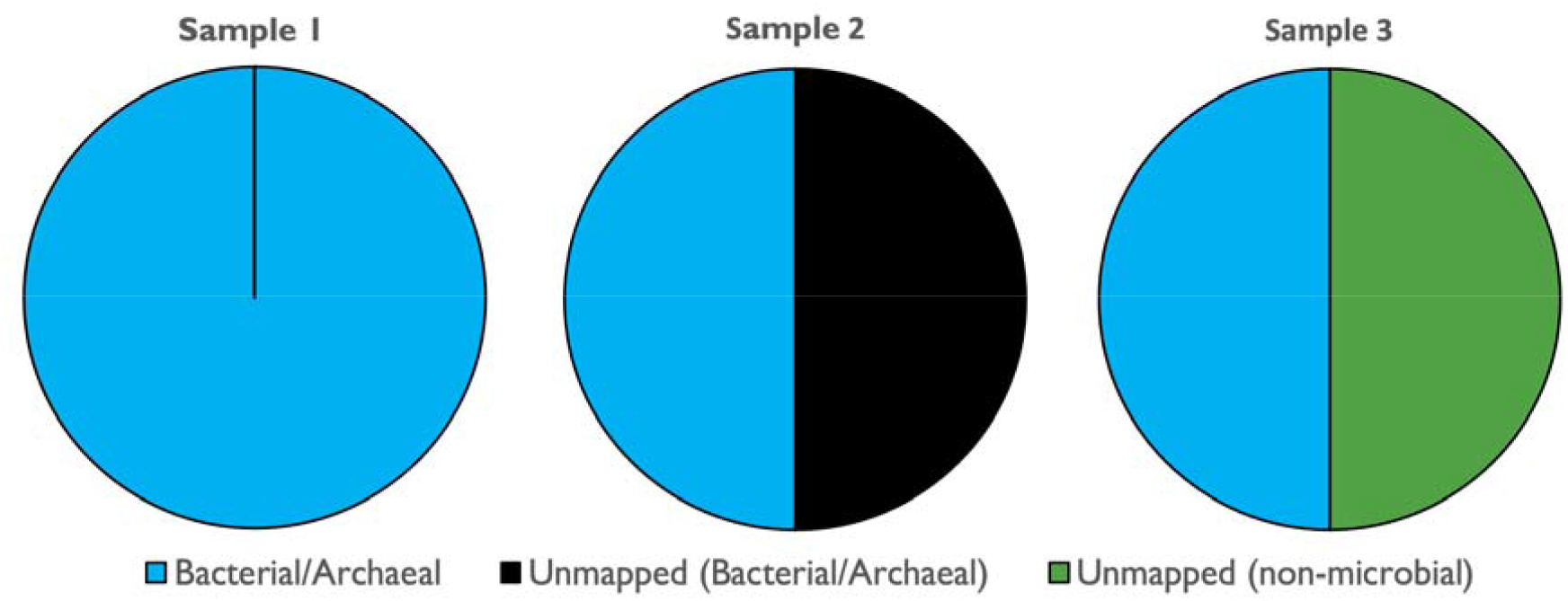
Non-microbial DNA obfuscates genome recovery appraisal. Metagenomic samples can have differing proportions of microbial DNA. In sample 1, all reads are microbial and are recruited to the MAG catalogue, suggesting successful genome recovery. In sample 2, all the reads are microbial but the MAG catalogue is not representative, resulting in only 50% of reads mapping. Sample 3 contains 50% microbial and 50% non-microbial DNA (from e.g. host or dietary sources), so even a perfect microbial genome recovery will only result in 50% of reads mapping. If non-microbial DNA is not removed, the success of genome recovery for samples 2 and 3 will be judged as equal by the rate of reads mapping, even though sample 3’s microbial community is far better represented.

When non-microbial sources of DNA are known *a priori*, and high-quality reference genomes of these sources are available, filtering the corresponding reads from metagenomes is straightforward. For example, human saliva samples typically have a high proportion of host reads (>80%), which can be removed by mapping to a human reference genome (11). Indeed, removal of human reads before submission to public sequencing databases is typically considered an ethical requirement for human metagenome studies (12, 13). However, in microbiomes from other hosts and environments, removal of non-microbial reads is often intractable. Even if a high quality host genome is available, there is a lack of reference genomes for most dietary species (e.g. plants, insects, etc.) and non-microbial species to be included in a read mapping database may not be known (e.g. novel fungal diversity in soil metagenomes) (14). Therefore, to properly inform genome recovery efforts, a method to accurately estimate the proportion of microbial reads in metagenomes is required. Estimation of this proportion (hereafter ‘microbial fraction’) would also guide researchers attempting to enrich, deplete, or maintain microbial DNA through laboratory-based cell size filtration or differential cell lysis methods (15, 16).

In this paper, we introduce ‘SingleM microbial fraction’ (hereafter ‘SMF’), a tool for estimating the quantity of bacterial and archaeal DNA in metagenomic datasets. Instead of filtering out non-microbial reads, SMF bases its estimate purely on the detection of reads encoding *microbial* single copy marker genes. An important advantage of this approach is that it does not require any knowledge of *non-microbial* genomes such as Eukaryota or viruses, which are often missing from reference genome databases. Accurate detection of entire microbial communities was recently made possible by the development of SingleM, a tool which reliably detects microbial species even when they are not represented in reference genome databases (17). We validate SMF on simple, complex, and diverse simulated metagenomes. Two real datasets are then used to demonstrate how SMF can improve evaluation of genome recovery efforts. Finally, we apply this fast and scalable approach to 136,284 publicly available metagenomes to demonstrate that variation in microbial fractions exists both within and between sample types.

## Methods

### Implementation of SingleM microbial fraction (SMF)

The SingleM microbial fraction (SMF) workflow is implemented as a post-processing step (implemented in the `singlem microbial_fraction` mode) of community profiles generated by the main workflow of SingleM (`singlem pipe`) (17). The main workflow of SingleM takes as input raw metagenomic reads and estimates a community profile, composed of Genome Taxonomy Database (GTDB) (18) taxons alongside their estimated per-base read coverage. This coverage is an approximation of the total read coverage of all genomes belonging to the taxon. For instance, the coverage of a species is the sum of the coverages of each strain in the metagenome.

The community profile output by SingleM provides the information required to estimate the relative abundance of each taxon. This can be calculated by dividing the coverage of each taxon by the sum of all coverages. However, the per-base read coverage can also be used to estimate the total abundance of reads in the metagenome belonging to that taxon. This is what SMF does.

At its core, SMF operates by applying a simple equation for each species similar to the Lander/Waterman equation (19):

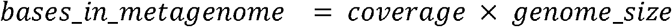

Where *coverage* is the average number of reads covering each base in the species’ genome. Here we assume a single *genome_size* for each species, and use SingleM’s estimate of the *coverage*. We then estimate the total number of bases in the metagenome that derive from all microbial genomes by summing the estimates for each species:

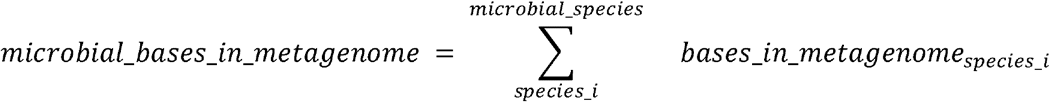

And finally estimate the fraction of the metagenome which is microbial, with a maximum of 100%:

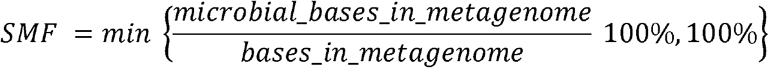

### Estimation of genome size for SMF calculation

To estimate the true *genome_size* of each species, accounting for imperfections which arise from genome recovery efforts, we take the genome size of the GTDB species representative and adjust it based on its completeness and contamination as predicted by CheckM2 (20):

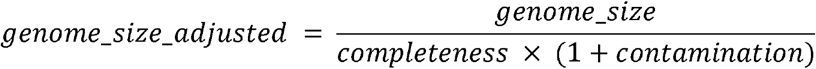

Community profiles generated by SingleM also contain entries from lineages not resolved to the species level e.g. coverage assigned to the genus or family level. For these cases we use genome sizes estimated from lower taxonomic ranks. For genera:

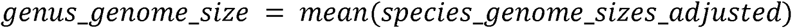

Where *species_genome_sizes_adjusted* is the set of adjusted genome sizes of species within the genus. For family, order, class, phylum and domain levels we average the average genome sizes of the taxons immediately below them. For families:

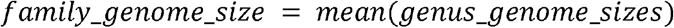

Where *genus_genome_sizes* is the set of genome sizes of genera belonging to that family. Orders, classes, phyla and domains are calculated similarly based on the mean genome sizes of their families, orders, classes and phyla respectively.

### Warnings about unreliability of estimates for some communities

Some communities are dominated by individual lineages which are not currently represented at the species level in reference genome databases. These communities are challenging for the approach used here because incorrect genome size estimation(s) can have a large impact on SMF. In simple communities composed of a few genomes, the SMF value can vary markedly with the (uncertain) estimate of only one or two lineages (**Supplementary Note 1**).

In more complex communities such as those found in soils and many animal guts, the genome sizes of individual lineages may also be uncertain. However, in these cases SMF is derived from many independent estimates of genome size (one for each lineage), even if the individual estimates are uncertain. For SMF to be inaccurate, many genome size estimates have to be incorrect, and either systematically overestimated or systematically underestimated. Therefore, we conclude that more complex communities are less affected by uncertain genome size estimates.

Following this observation, `singlem microbial_fraction` emits a warning when simple communities with novel species are encountered. Specifically, we consider the three most abundant lineages not classified to the species level in the profile. If doubling or halving their estimated genome size changes the SMF by more than 2%, then a warning is emitted.

Users can improve the accuracy of SMF by enriching the SingleM reference database with user-generated MAGs, i.e. producing an updated SingleM ‘metapackage’. This can be achieved by using `singlem supplement` to add the new MAGs, calculate community profiles using this metapackage with `singlem pipe` and then use `singlem microbial_fraction` to generate updated SMF values. An example of this workflow is provided in **Supplementary Note 1**.

### Estimation of average microbial genome size in a metagenome

Average microbial genome size in a metagenome is calculated by `singlem microbial_fraction` using the per-taxon genome sizes estimated according to the procedure above. The average genome size is the average of these genome sizes after weighting them by their coverage in the SingleM community profile. That is, it is calculated according to:

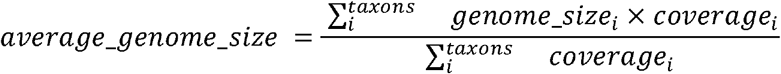

The approach used here differs from previous methods at estimating average genome size which assume that the entire metagenome is derived from microbial genomes(21, 22).

### Benchmarking using simulated data

We benchmarked the performance of SMF on four simulated datasets. Gen-paired-end-reads (https://github.com/EisenRa/reads-for-assembly) [forked from merenlab/reads-for-assembly] was used to generate simulated paired-end reads from the Zymo Mock Community Standard reference genome files (ZymoBIOMICS.STD.refseq.v2; also available on our GitHub repository), host reference genomes (*Homo sapiens*: GCF_000001405.39_GRCh38.p13; *Arabidopsis thaliana*: GCF_000001735.4_TAIR10.1; *Plasmodium falciparum*: GCF_000002765.5_GCA_000002765), and CAMI II marine and strain-madness reference genomes (profile #0 for each CAMI II dataset) (23). For more information and reproducible code, see (https://github.com/EisenRa/SingleM_microbial_fraction_paper).

In a separate simulation to benchmark the performance of SMF on communities including divergent species (**Figure 2C**), we used a Snakemake (24) workflow available at https://github.com/wwood/singlem-read-fraction-benchmarking. Reads were simulated for 120 genomes from species which were added to the GTDB database in R214. Reads were simulated using ART (25) at 10X coverage of two microbial genomes, one novel, the other present in GTDB R207. The pair of genomes in each mock community was chosen to have similar genome sizes and such that one was bacterial and the other archaeal. SingleM v0.16.0 was run first in “pipe” mode using the S3.1.0 metapackage (doi:10.5281/zenodo.7582579) to generate a GTDB R207 community profile from the simulated reads. The `microbial_fraction` mode was used to apply SMF using the same metapackage but an updated taxon-genome-lengths file available in the benchmarking code GitHub repository above (https://github.com/wwood/singlem-read-fraction-benchmarking).

**Figure 2:**
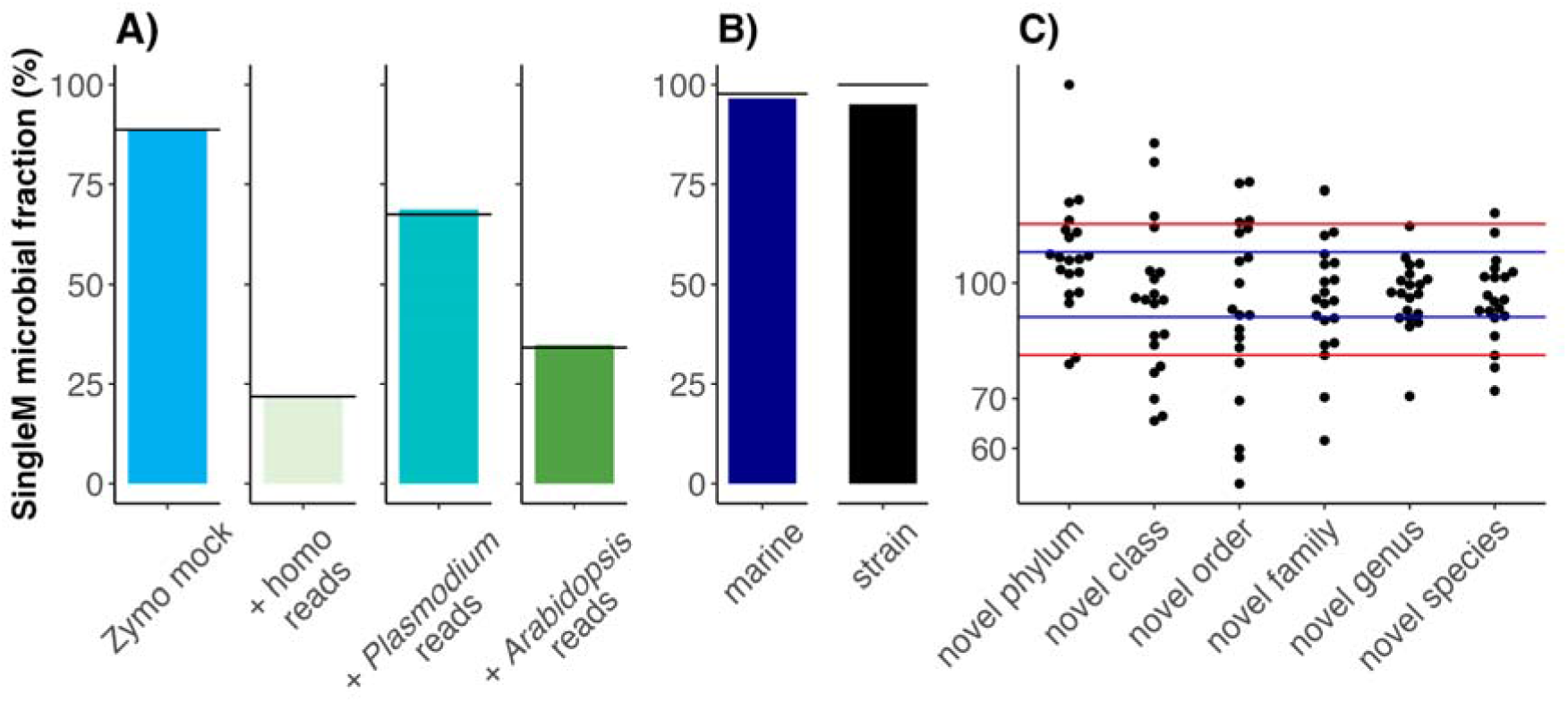
Performance of SMF on simulated metagenomes. **A)** Simulated eukaryotic reads from *Homo sapiens, Arabidopsis thaliana*, and *Plasmodium falciparum*, were spiked into simulated Zymo mock community metagenomes. Horizontal bars represent the true proportion of bacterial and archaeal DNA in the simulated metagenomes. **B)** Simulated reads based on community compositions from the more complex CAMI II marine and strain-madness datasets. **C)** Tests on metagenomes containing divergent genomes. Reads were simulated from 120 mock communities (dots) which were each composed of 2 species at equal abundance, one present in GTDB R207 and a second which was new in GTDB R214 at varying degrees of novelty. SMF was run on each community using an R207 reference database. Blue and red lines represent 10% and 20% error respectively from the true answer of 100%. Given the challenging nature of estimating non-microbial DNA fractions in these communities, for many of these communities a warning message appears indicating that the estimate may be unreliable (**Supplementary Figure 1**).

### Analysis of previously published data

Sequencing reads from human (26) and spotted hyena (27) faecal samples were downloaded using Kingfisher (28). Ten random human faecal samples were selected, and randomly subsampled to 5 Gbp of data using seqtk sample (https://github.com/lh3/seqtk). Hyena samples from four individuals (8 samples per individual) were used, and host reads were removed by mapping to the spotted hyena reference genome (GCA_008692635.1) using BowTie2 (29). SingleM version 0.16.0 was used to estimate microbial fractions with metapackage 3.2.1 (https://doi.org/10.5281/zenodo.8419620), which was based on GTDB (18) R214. For both datasets, the Earth Hologenome Initiative (EHI) genome-resolved metagenomic pipeline was used (https://github.com/earthhologenome/EHI_bioinformatics). Briefly, reads were quality trimmed using fastp (30) and mapped to their respective host genomes using Bowtie2 (29), before removing mapped reads with samtools (31). Unmapped reads were then coassembled using megahit (6), before being binned and refined using MetaWRAP’s binning (MetaBAT2 (32), MaxBin2 (8), CONCOCT (7)) and refinement modules (33, 34). The resulting bins were dereplicated at 98% ANI using dRep (35), and the processed reads were then mapped to their respective MAG catalogues using Bowtie2. Final count tables were created with CoverM (36) using --min-covered-fraction 0.

### SMF estimates across public metagenome datasets

To calculate the SMF across public metagenomes, taxonomic profiles from Sandpiper 0.2.0 were taken as input to “singlem microbial_fraction” v0.16.0 using the --input-metagenomes option, supplying metagenome sizes derived from the NCBI SRA BigQuery SQL database (column “mbases”). Samples were filtered by SRA metadata: library_strategy == “WGS” & library_selection == “RANDOM”. Only samples with > 0.5 Gbp were analysed. As the SRA can contain samples with incorrect metadata entries, for some analyses, we opted for the usage of samples that had been manually curated. We used samples with curated metadata from the AnimalAssociatedMetagenomeDB (37), MarineMetagenomeDB (38) for figures 4D/E, respectively. Curated location metadata from (39) was used for human faecal samples in figures 4 B/C.

For soil metagenomes, metagenomes with very low SMF values (<5%) were excluded from analysis as manual inspection of a random sample of these showed that these samples were not standard metagenomes e.g. they were single cell amplifications or targeted at mitochondria. Metagenomes which had no annotated latitude or longitude were also excluded.

### Estimating DAMR for marine metagenomes

Mapping rates to different marine genome catalogues were obtained from Nishimura et al. (supplementary file 1) (40).

### Estimation of fungi:microbial cell ratios in soil metagenomes

To convert an SMF value, a percentage of DNA reads that are microbial, into an estimate of the relative abundance of fungal and microbial cells, the following simplifying assumptions were made:

1. The community is largely composed of fungi and bacteria only, with the remainder contributing very few reads.
2. The metagenomic data are derived from DNA extraction methods which lysed fungal and bacterial cells with equal efficiency.
3. The abundance-weighted average size of fungal genomes in each sample is 37.5 Mbp. This average was derived by taking the median genome size of each Ascomycota (41) genus, and then taking the mean of those median values.

The average genome size of microbial community members was estimated for each metagenome with the method described above. To calculate cellular ratio from the

SMF, we used the following formula, where *c*_*m*_ is the number of microbial cells, *c*_*f*_ is the number of fungal cells, *g*_*m*_ is the average microbial genome size, and *g*_*f*_ is the average fungal genome size.

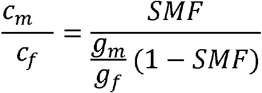

This equation was derived starting from the following, where *r*_*m*_ is the number of reads derived from microbial cells, *r*_*f*_ is the number of reads derived from fungal cells, and *s* is the average number of reads generated that start at each genomic position:

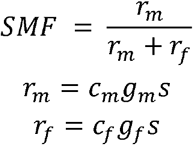

Taking the ratio of the last pair of equations, multiplying it by *r*_*f*_ and substituting into the first equation leads to the formula above for 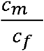 .

## Results and discussion

### Benchmarking on simulated metagenomes

We first tested SingleM microbial fraction (SMF) on a simple simulated metagenome, composed of 8 bacterial and 2 fungal species (“Zymo Mock”: based on the ZymoBIOMICS™ Microbial Community Standard). SMF estimated the microbial fraction to be 88.80%, very close to the 88.68% ground truth (**Figure 2A**). The estimated microbial fraction was also within 1% of the correct value when simulating metagenomes with varying proportions of spiked-in eukaryotic DNA (**Figure 2A**).

Next, we tested SMF on more complex simulated metagenomes: A “marine” benchmark dataset based on compositional profiles of the deep sea environment from the CAMI II challenge (23), consisting of 389 bacterial and archaeal species (and 200 plasmid/viral sequences); and a “strain-madness” benchmark dataset comprising 405 bacterial and archaeal genomes with high strain-diversity (97% of genomes have a closely related strain). SMF estimated 94.3% and 95.1% of the reads to be microbial, respectively, close to the true microbial fractions of 97.7 and 100% for these communities (**Figure 2B**). These results suggest that SMF is not adversely affected by high community diversity.

In the final and most challenging benchmark, we tested the performance of SMF under worst-case scenarios, when communities containing novel lineages were present (**Figure 2C**). Simulated communities contained two microbial species at equal abundance (10X coverage each), one present in both R207 and R214 releases of GTDB, and another only in R214. In each simulated dataset, the new genome had a different level of taxonomic novelty (from species to phylum) relative to R207. These metagenomes were analysed using SMF backed by the GTDB R207 database. SMF estimates were within 20% of the true value of 100% for most communities containing novel species (80%), genera (95%) and families (85%), but was less accurate for novel orders (55%), classes (60%) and phyla (70%). We consider this final set of benchmarks especially challenging since the communities contained very high quantities of very novel lineages. Communities dominated by species novel at the order level are increasingly rare as genome databases move towards completion, and in these situations combining SMF with a genome-centric approach might be appropriate (**Supplementary Note 1**). Recognising this situation, SMF emits a warning when simple communities containing novel lineages are encountered, a situation we found to be rare in practice (**Supplementary Figure 1, Supplementary Note 1**).

### Benchmarking on real metagenomes

We next tested SMF on real metagenomic data to demonstrate how it can be used to evaluate genome recovery efforts.

#### Human faecal samples

Human faecal metagenomes usually contain ∼90% microbial DNA (42), and can therefore be used as a control to benchmark SMF. After subsampling sequencing data sets to 5Gbp per sample, genomes from 10 faecal samples (26) were recovered yielding 200 medium or high quality MAGs. The fraction of reads mapped against these MAGs was 46.6% ± 4.1% (**Figure 3A**), much less than expected. In contrast, SMF estimates of close to 100% (96.7 ± 3.23%) agreed with the *a priori* expectation. The SMF estimates therefore predicted that the low mapping rates were due to challenges in genome recovery rather than the presence of non-microbial DNA in the metagenomes. This prediction was confirmed by mapping reads to the more extensive Human Reference Gut Microbiome genome catalogue (26), which recruited 90.4 ± 2.36% of reads. These results show that SMF can be used to attribute poor mapping rates to imperfect genome recovery when metagenomes are dominated by microbial reads.

**Figure 3:**
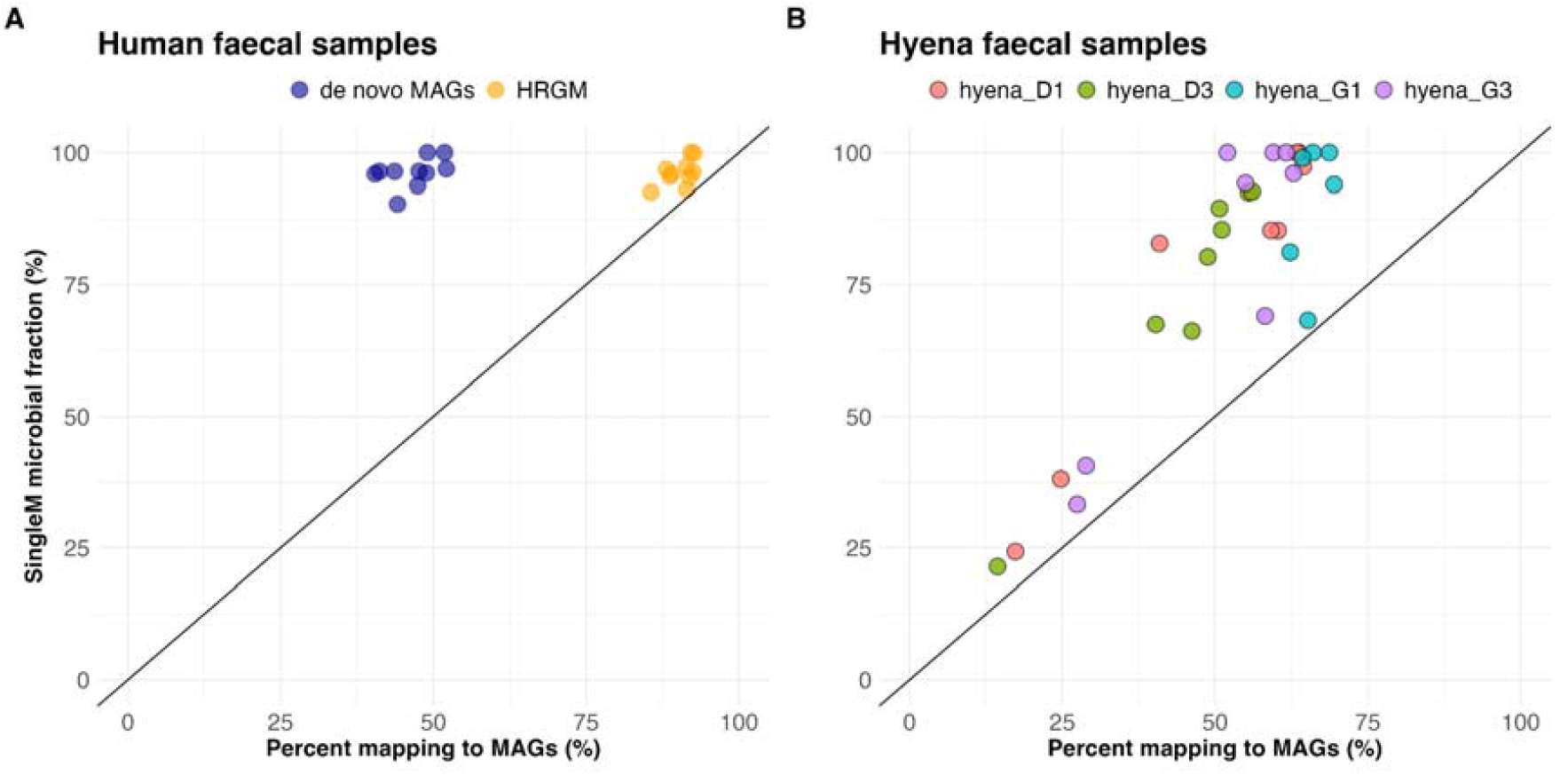
SingleM microbial fraction (SMF) can help evaluate genome recovery efforts. **A)** Human faecal dataset. Percentage of reads mapping to the *de novo* MAGs (blue) versus the percentage of the metagenomes estimated to be microbial by SMF. Here the *de novo* dereplicated MAGs underrepresent the true diversity in the samples, whereas most reads are recruited to the Human Reference Gut Microbiome (HRGM: yellow). **B)** Hyena faecal dataset. Percentage of reads mapping to the MAG catalogue versus SingleM microbial fraction estimates. Points on or close to the straight line indicate that the de novo MAGs were sufficient to capture most of the diversity in the sample.

#### Hyena faecal samples

Unlike humans, non-model species often lack microbial or host reference genomes, and may contain varying quantities of DNA derived from dietary and/or protozoan sources (43, 44). Hyenas are carnivores with diets that can fluctuate throughout the year, so we expected that faecal samples could contain varying proportions of prey DNA from their diets. This variability can make it difficult to determine whether the microbial diversity in a sample is adequately captured using standard assessments. We applied SMF to a recent study on wild spotted hyenas (*Crocuta crocuta*)—the first metagenomic study for this species (27). Each hyena was sampled multiple times, so we coassembled metagenomic reads from hyena faecal samples by individual, and binned contigs into *de novo* MAGs. Our reprocessing improved the mappability of each sample’s reads to the *de novo* MAGs substantially compared to the original study (original study: 13.2 ± 7.7%, here: 52.8 ± 15.5% SD). SMF estimated substantial variation in the proportions of microbial DNA for these samples (20.2% — 100%) (**Figure 3B**).

### Domain-adjusted mapping rate (DAMR)

Application of SMF to these human and hyena faceal samples demonstrates that mapping rates alone can be a very poor estimation of MAG recovery success when the number of microbial reads is unknown. To overcome this limitation, here we introduce a new metric named “Domain-Adjusted Mapping Rate” (DAMR). DAMR is calculated as the rate of read mapping to sets of microbial genomes (MAGs or other references) divided by the fraction of the community predicted to be bacterial or archaeal (here with SMF). For example, if a metagenome is predicted to have 50% microbial DNA, and 45% of reads map to the MAG catalogue/references, the DAMR is 90% (45% / 50%). In rare circumstances where the SMF estimate is lower than the fraction of mapped reads, indicating artefacts in mapping and/or SMF, then the DAMR is rounded down to 100%. In the human analysis outlined above, the DAMR of the Human Reference Gut Microbiome genome catalogue was calculated to be 93.5%. In the hyena example, combining SMF estimates with the proportion of reads mapping to the *de novo* MAGs yielded DAMR values ranging from 49.3% — 95.6%, which allowed us to identify samples that may require additional sequencing to improve genome recovery efforts. In summary, combining SMF with the DAMR metric can be a valuable tool for interpreting the performance of prokaryotic genome recovery from understudied host species and complex metagenomic mixtures. This synergy facilitates the enhancement of data quality by pinpointing samples requiring further characterisation through supplementary sequencing or bioinformatic efforts.

### Microbial read fractions of Earth’s metagenomes

To explore global trends in microbial fractions amongst samples sourced from a broad range of environments, we applied SMF to publicly available metagenomic datasets. In total, SMF was calculated from 136,284 community profiles available on the Sandpiper (17) website. These estimates were subsequently incorporated back into Sandpiper, where SMF values are now shown for each metagenome.

SMF estimates were first compared to those produced by STAT (45), a kmer-based tool for taxonomic assignment of reads. The tool is trained on RefSeq genomes and is widely available for metagenomes at NCBI SRA. We hypothesised that because STAT is reliant on matching kmers to reference genome data, SMF would provide better estimates for understudied sample types or host species. SMF and STAT estimates were within ±5% of each other for 18.2% of samples. SMF gave higher microbial fraction estimates (at least >=5%) than STAT for 58.6% of metagenomes (**Figure 4A**). In 33.2% of samples, the SMF estimation was >2-fold higher than STAT, consistent with our hypothesis that STAT underestimates metagenomes containing novel lineages. Conversely, in only 7.1% of metagenomes was STAT >2-fold higher than SMF (28.5% of these were ‘viral metagenomes’).

**Figure 4:**
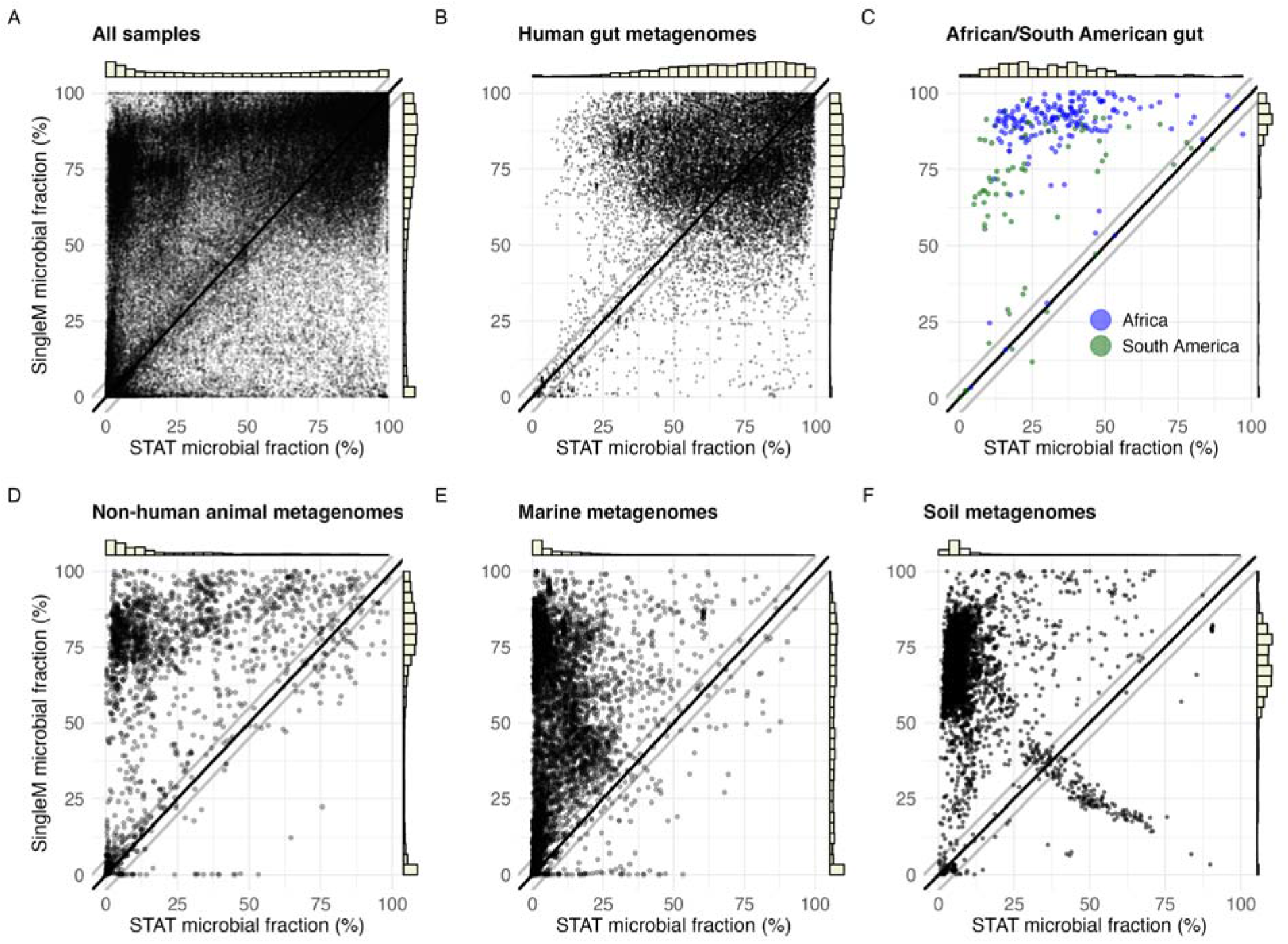
Comparison of microbial fractions derived from STAT and SingleM microbial_fraction. SingleM microbial fraction (Y-axes) was run on 136,284 metagenomes from the SRA and compared to STAT microbial fraction estimates (X-axes). **A)** All samples. **B)** 18,571 metadata-curated human gut metagenomes. **C)** African and South American human faecal samples. **D)** 2,073 curated, non-human samples. **E)** 5,490 curated marine metagenome samples. **F)** 4,160 soil metagenome samples. Black lines represent equal SMF and STAT values, grey lines represent ± 5%. The bar charts represent the distributions of SMF and STAT values. The small cluster of samples with a higher STAT estimate than SMF below the diagonal are discussed in **Supplementary Note 2**.

We next investigated samples from different environments. For human faecal metagenomes we restricted the analysis to metagenomes with location information curated by Martiny et. al. (39) (n=18,571, **Figure 4B, Figure 5A**). As expected, human gut metagenomes typically showed a high microbial fraction (mode ∼ 90%) (**Figure 5A**). SMF estimates were somewhat higher than STAT (SMF/STAT: median 77.2%/69.7%, SD 18.5%/21.3%), suggesting that most human faecal samples are well represented by reference databases. However, SMF estimates were substantially higher than STAT in samples derived from understudied human populations: African (median 91.8% vs 34.0%, Mann-Whitney *p*-value 3.68e-51) and South American (median 74.9% vs 21.4%, Mann-Whitney *p*-value 2.21e-15) populations (**Figure 4C**). One plausible explanation is the underrepresentation of African and South American gut microbial genomes in the public databases. Most work on the human gut microbiome to date has been from individuals from the Global North (∼71% of human faecal samples are from Europe, U.S.A, or Canada (46)).

**Figure 5:**
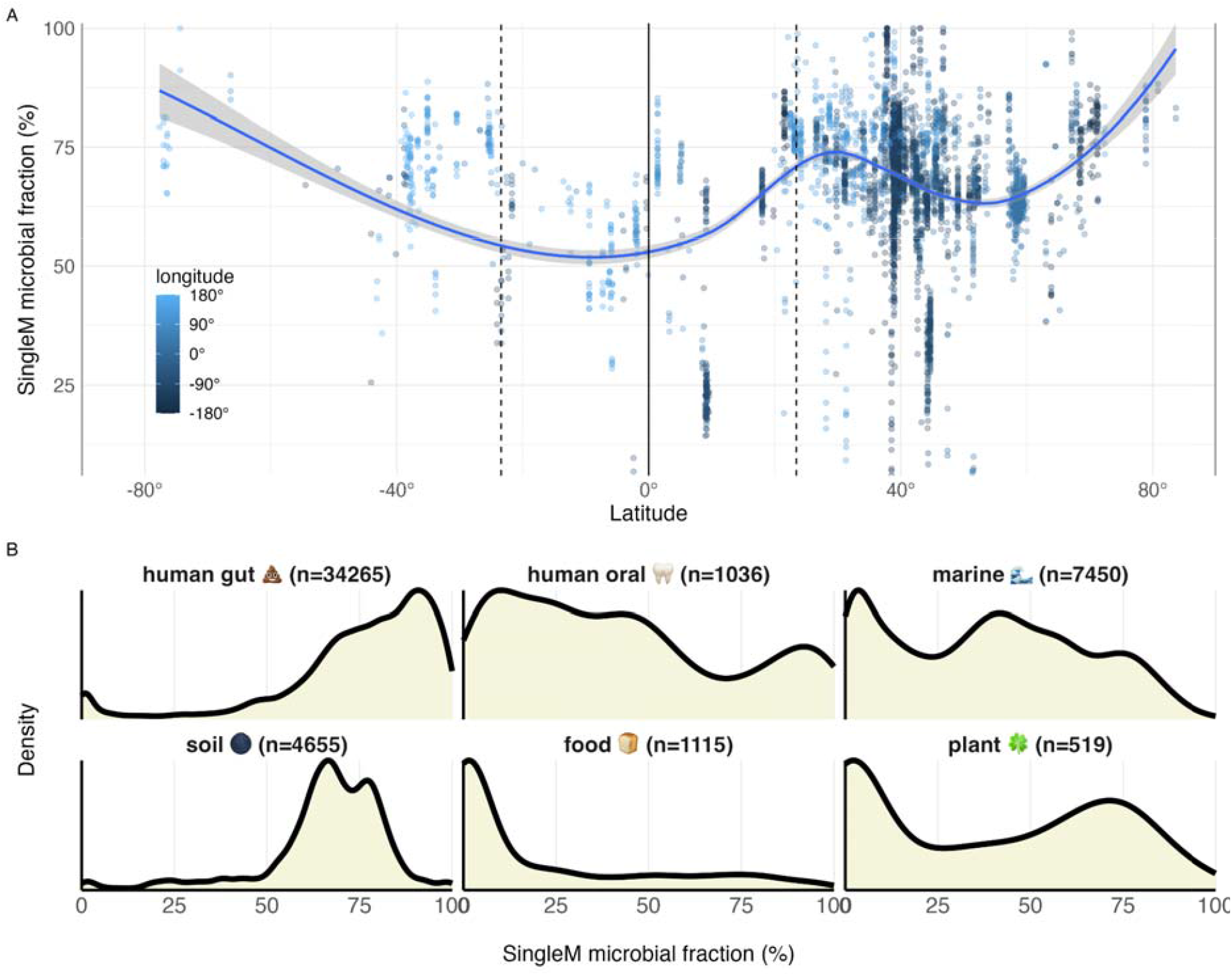
SingleM microbial fractions of metagenome sample types sourced from the SRA. **A)** Association between soil metagenome microbial fraction and latitude/longitude. **B)** Density plots of SingleM microbial fractions grouped by sample type.

Genomes derived from non-human animal microbiomes are also considerably underrepresented in public databases. To explore this, we used the set of SRA samples curated from the AnimalAssociatedMetagenomeDB (37) (n=2,073 samples). SMF estimates were again higher for these samples (median = 76.5 SD = 31%) compared to STAT (median = 12.4 SD = 23.9%, **Figure 4D**), consistent with the hypothesis that SMF is more capable of estimating microbial fractions from metagenomic samples of understudied hosts. This was also the case for the subset of better-studied domestic animals (Chicken n=352: SMF median 89.5%, SD 12.7%; STAT median 42.9%, SD 19.6% & Bovine n=202: SMF median 78.4%, SD 22.1%; STAT median 7.4%, SD 5.6%).

Marine metagenomes are derived from the largest biome on Earth, with samples sourced from varying geographic locations, depths and sample types (e.g. water, sediment). SMF yielded much higher microbial fraction estimates than STAT (**Figure 4E**) as expected, given the paucity of RefSeq quality reference genomes from marine environments (40). Marine water metagenomes are typically generated by filtering seawater through pores of a defined size (16). The size of these pore varies between studies, where ultrasmall (<0.22μm) size selection targets viruses, intermediate (0.22-3μm) sizes target Archaea and Bacteria, and larger sizes (3μm+ or 20μm+) enrich for eukaryotic plankton (16). Marine metagenome SMFs were strongly influenced by filtration size, with ultrasmall (17.7 ± 15.0%) and larger sizes (3.7 ± 6.1%) exhibiting a much smaller SMF than intermediate sizes (70.6 ± 13%, **Supplementary Figure 3**). We also used SMF values to assess the performance of recent efforts to create a unified marine genome reference catalogue for marine metagenomes (40). The estimated DAMR for the Unified Genome Catalog of Marine Prokaryotes (UGCMP) was 63.4% (SD 13.6%) (**Supplementary Figure 4)**, suggesting that many marine microbial genomes are yet to be recovered.

Soil metagenomes were found to contain mostly microbial DNA, with 88% having SMF >50% (median 69.0%, SD 15.1%, **Figure X5**). Soil microbial communities are poorly represented in genome databases (17, 47), so STAT estimates were almost always much lower (14% had an estimated microbial fraction >50%, and 68% were <10%, **Figure 4F, Supplementary Note 2**). These analyses confirm SMF’s ability to estimate microbial fraction in metagenomes containing many novel species. They also show that the well known challenges for recovering microbial genomes from soil (47–49) are not usually a result of eukaryotic DNA “contaminating” these metagenomes.

Soils harbour many different kinds of organisms, with bacteria and fungi thought to be the most dominant members overall. The ratio of fungi to bacteria, however, can be variable. Using quantitative amplicon-based sequencing, Fierer et al. observed ratios of 0.007 to 0.34 amongst a range of unmanaged soil environments (50). Under some simplifying assumptions, the median SMF value (69.0%) corresponds to a fungal to microbial cell ratio of 0.053 (see methods). The 90th and 10th SMF percentiles for soil (82% and 48%) correspond to fungi:bacteria cellular ratios of 0.03 and 0.13, in rough agreement with Fierer et. al. (50).

Many publicly available soil metagenomes are annotated with the GPS coordinates of their sampling location. Latitude was related to SMF non-linearly, with significantly less microbial DNA in metagenomes sampled between the tropics of cancer and capricorn (median 63.0%, SD 13.6%) compared to those outside these parallels (median 70.0%, SD 13.6%, p-value =4e-59, t-test) (**Figure 5A**). Assuming that fungi are the major source of non-microbial DNA, these results suggest that soil communities in the tropics may have a higher fungi:microbial ratio than those outside this region. Eukaryote:microbial cell ratios are no doubt driven by patterns more complex than those described by latitude alone (e.g. C:N ratio (50)or pH (51)), and we hope SMF will be used in future to help unravel these mysteries.

Overall, metagenomes derived from different environments had substantially different microbial fractions, and individual environment types were often heterogeneous (**Figure 5B**). For instance, samples labelled ‘human oral metagenome’ had a wider distribution of microbial fractions, which likely relates to the diversity of distinct oral niches (**Figure 5B**) (52). Saliva samples typically have high proportions of host DNA (>80%), whereas tongue swab samples have lower proportions (∼10%) (42). Most of the ‘food metagenome’ samples had low microbial read fractions (median 9.2%), highlighting potential issues inherent with these sample types, such as high quantities of raw ingredients. For instance, we found cow DNA in milk SRA (run ERR3143475 (53) had SMF 1% and 76% of reads assigned to the family Bovinae according to STAT) and likely fungal DNA in ferments (SRR648391, a metagenome derived from soy sauce, had SMF 24%). Plant metagenomes exhibited a distribution of different microbial fractions, likely related to variation in the types of samples in these categories (e.g. phyllosphere, rhizosphere) (40, 54). These results highlight that microbial fractions can differ both within and between biomes.

Raw metagenomes derived from human microbiomes contain substantial quantities of human DNA, particularly from non-faecal samples. However, since making human genetic data public might result in invasions of privacy, it is common practice to screen out human host DNA before submitting the data to publicly available repositories (55). We were therefore surprised to find many human microbiome samples with low SMF values (e.g. in human oral metagenomes, **Figure 5B**). After confirming many of these samples did indeed contain human DNA using STAT, we were forced to conclude that public metagenome databases contain large amounts of human genetic data. While metagenomic samples are usually deidentified, the availability of this data could have ethical and practical implications in some cases (13, 56).

## Conclusion

SMF can be utilised as a scalable and assembly-free means to guide researchers’ decision-making through prioritising samples prior to deeper sequencing, assessing microbial community representativeness, and benchmarking bioinformatic genome-recovery pipelines and laboratory method development. It will be especially useful as the field continues to explore novel sample types and hosts with unknown fractions of microbial DNA. Finally, the microbial read fractions generated for a large number of public metagenomes provides a new fundamental variable that can be used to explore how experimental or environmental variables shape microbial communities.

## Data availability

SingleM is available at https://github.com/wwood/singlem. Code and data for reproducing the analyses and figures of this paper are available at https://github.com/EisenRa/SingleM_microbial_fraction_paper and https://github.com/wwood/singlem-read-fraction-benchmarking.

## Acknowledgements

We thank Simon Roux and Matthew Sullivan for helpful discussion on marine datasets, and Joshua Mitchell for helpful suggestions for interpreting marine filter size metadata. B.J.W. was supported by Australian Research Council Future Fellow (#FT210100521) and Discovery Project (#DP230101171) grants. A.A. acknowledges the Danish National Research Foundation award DNRF143 ‘A Center for Evolutionary Hologenomics’ and the Carlsberg Foundation grant CF20-0460.

