## Supplementary information for "Large-scale estimation of bacterial and archaeal DNA prevalence in metagenomes reveals biome-specific patterns"

### Supplementary Notes

#### Supplementary Note 1

Seven hyena faecal samples yielded nonsensical SMF estimates over 100%. We found this was due to the actual genome sizes of one or a handful of species with high relative abundance (*i.e.* a highly uneven community) being much smaller than the mean genome size of their taxonomy in the GTDB. For example, sample G3_P301 had a highly abundant member classified by SingleM as g__Bacteroides, of which the mean genome size from the GTDB was 5.1 Mbp. The actual MAG size for this Bacteroides genome was 2.2 Mbp, resulting in >2.3 fold overestimation of its read contribution. We consider these scenarios to be rare, but have implemented a warning for users when a sample's SMF estimate could under- or overestimate the read fraction by >10%. Users are warned if the 3 highest abundance lineages not classified to the species level would change the estimated read fraction of the sample by >2% if their genome size is halved or doubled. In these situations, users should explore the samples that yielded warnings, and can, for example, update the SingleM estimates using genome sizes from MAGs that they generate to obtain more accurate estimates. In fact, when updating the estimates with MAG sizes from the hyena dataset, overestimations were drastically reduced (**Supplementary Figure 2**).

The most likely metagenomes to give rise to these situations are those dominated by a small number of highly abundant species. While it is challenging to estimate the microbial read fraction in these samples, MAG recovery from these samples is typically more successful since there is sufficient coverage and a comparatively less diverse community. The SMF algorithm warned about potential inaccuracies arising from species which were both highly dominant and novel in only 0.28% of public metagenomes, showing such cases are relatively uncommon in practice.

#### Supplementary Note 2

A small minority of soil samples had unusually high STAT values and low SMF values (200 samples, 4.8%, **Figure 4F**). Upon further investigation, it was found that 85% of these samples were derived from two studies that used the same tagmentation library preparation protocol. We downloaded the sequencing data for these bioprojects to estimate the insert sizes, and found a median insert size of 92 (± 23.3), suggesting that the library preparations in the original studies were not optimised (important for tagmentation-based protocols) (**Supplementary Figure 5**). Since SingleM relies on 20 amino acid (60 nucleotide) sequences, and these sequences must be contained within stretches of at least 72 bases uninterrupted by stop codons, libraries with short insert sizes would deflate SingleM microbial fraction estimates. For more details on the analysis and reproducible code, see <https://github.com/EisenRa/SingleM_microbial_fraction_paper/blob/main/code/SI_note2.md>

### Supplementary Figures

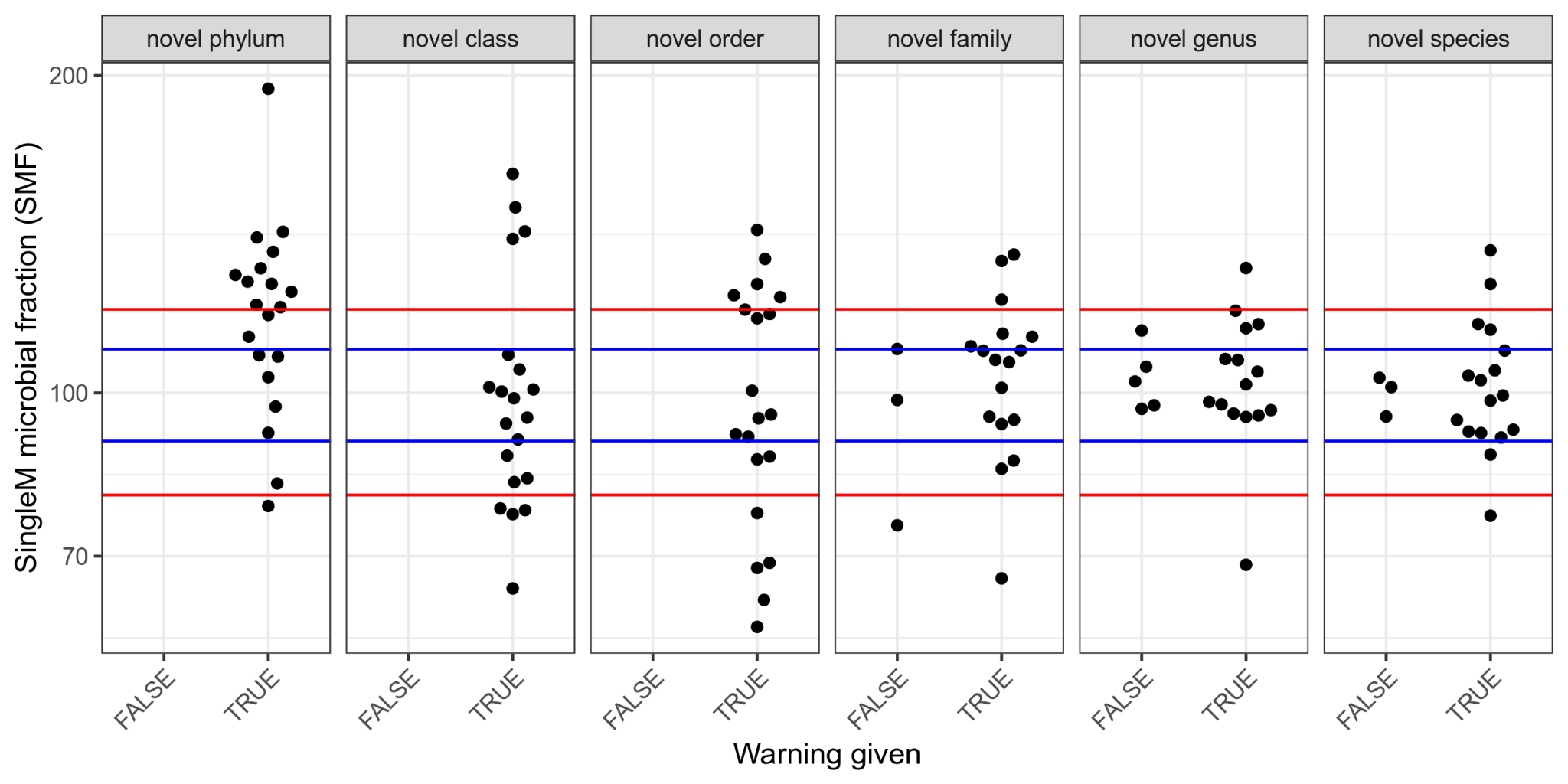

**Supplementary Figure 1**. Two-component community accuracy and whether a warning of an unreliable estimate is provided by SMF. This figure is the same as presented in **Figure 2C**, except that it also shows whether a warning is emitted by SMF.

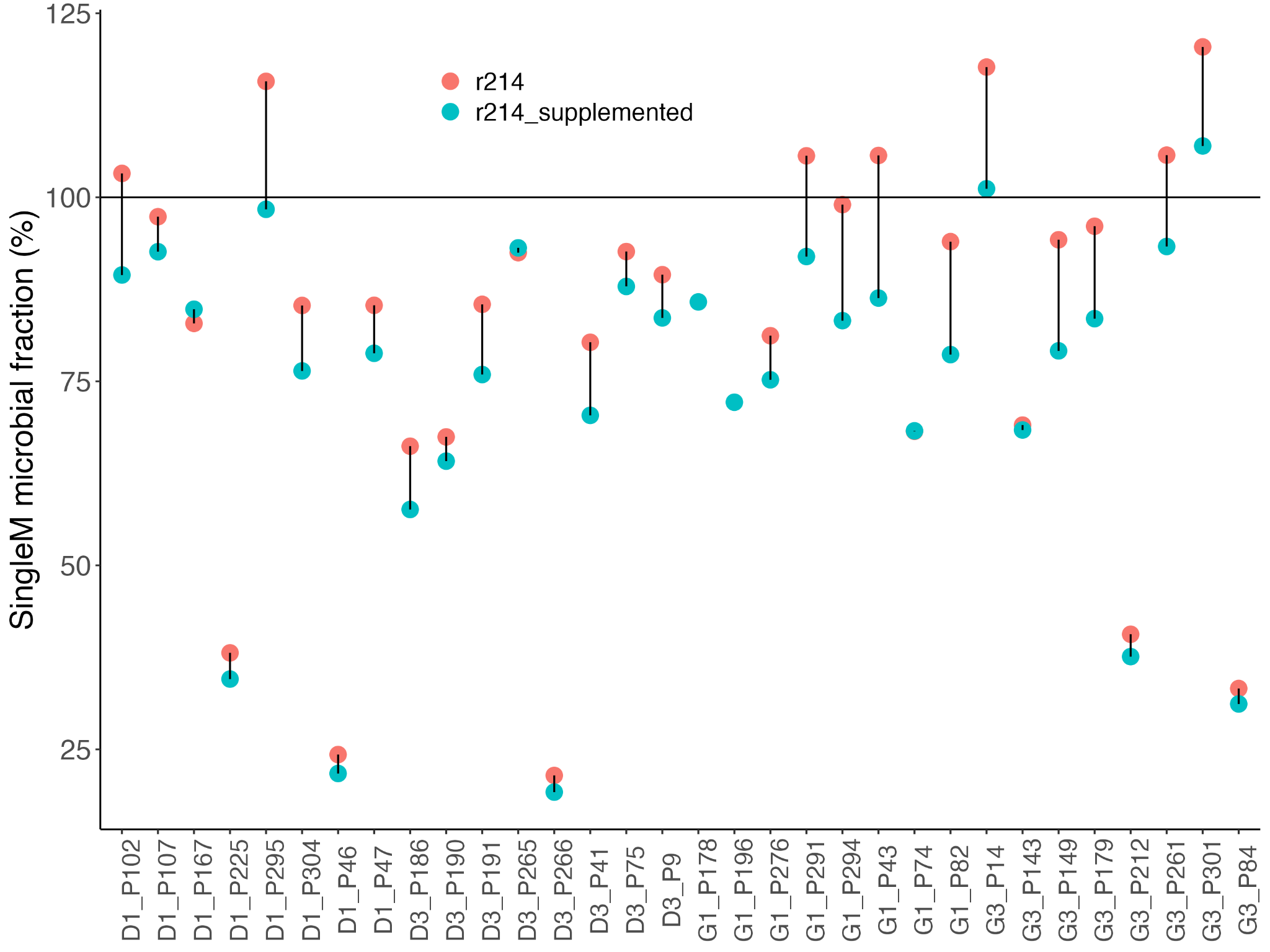

**Supplementary Figure 2**. SingleM microbial fractions of hyena faecal samples with default GTDB r214 genome sizes (red), and genome sizes supplemented from MAGs (blue).

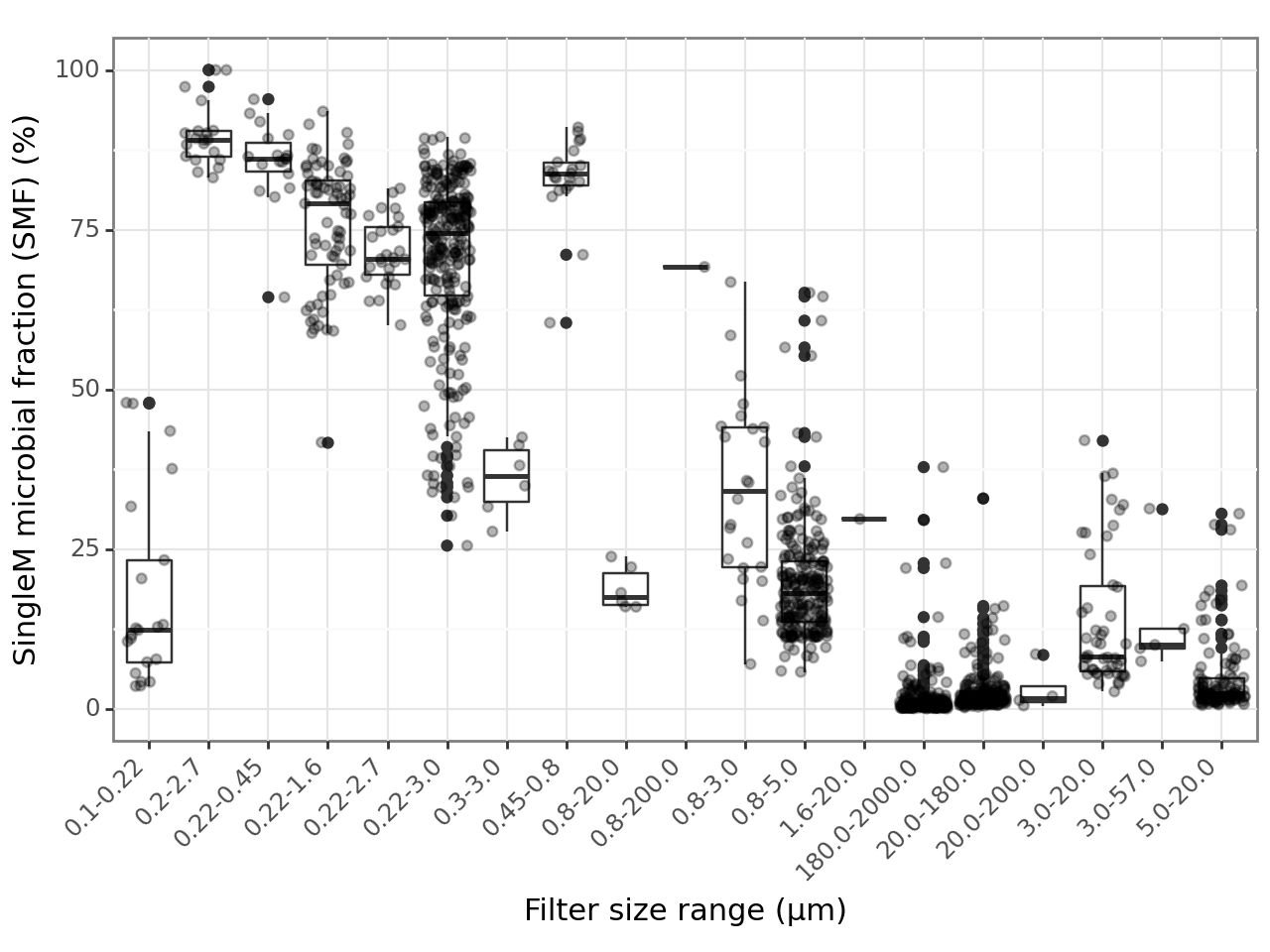

**Supplementary Figure 3. Size fractionation influences microbial read fraction in marine metagenomes.** Sequential size filtration is commonly used to separate cells of different sizes in the analysis of marine metagenomes. Where available as part of biosample metadata (Sunagawa et al. 2015; Pascoal et al. 2023; Hevroni et al. 2020; Hugerth et al. 2015), filter sizes (upper and lower sizes) were parsed and used here to stratify SingleM microbial fraction (SMF) estimates in marine metagenomes. SMF estimates were substantially affected by filter size, where the application of intermediate size bands between 0.22 and 0.8 μm resulted in higher SMF values, compared to smaller and larger filter sizes. This observation is consistent with established practice, since these smaller and larger sizes are thought to enrich for viral and eukaryotic cells, respectively, while intermediate sizes are thought to enrich for Bacteria and Archaea.

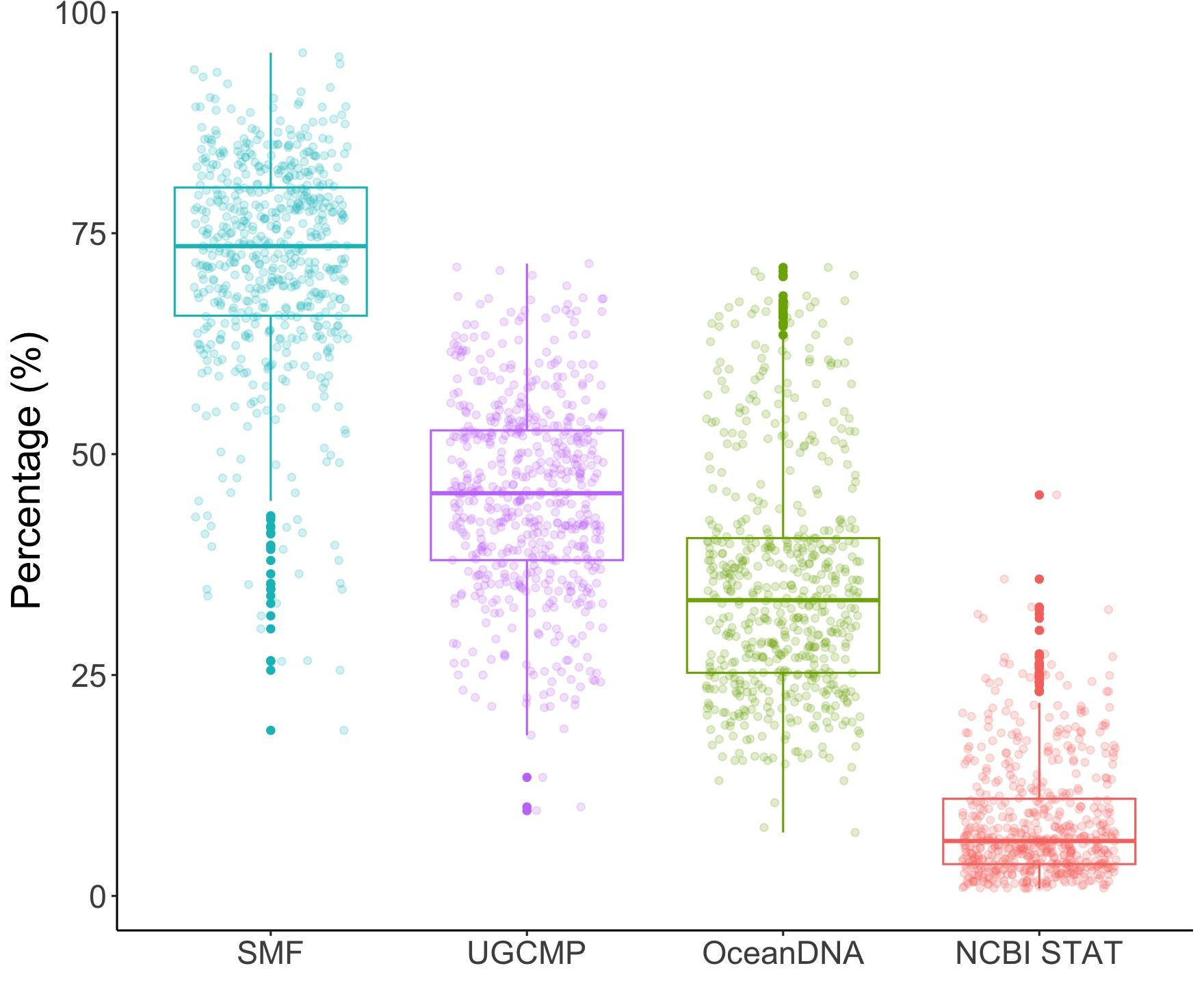

**Supplementary Figure 4**. Use of SMF to estimate the representativeness of marine genome catalogues. SMF = SMF estimate of metagenome. UGCMP = mapping rate of samples to the Unified Genome Catalog of Marine Prokaryotes. OceanDNA = mapping rate of samples to the OceanDNA MAG catalogue. STAT = STAT estimate of metagenome microbial fraction. Mapping rates were obtained from Nishimura et al. 2022 supplementary table 1.

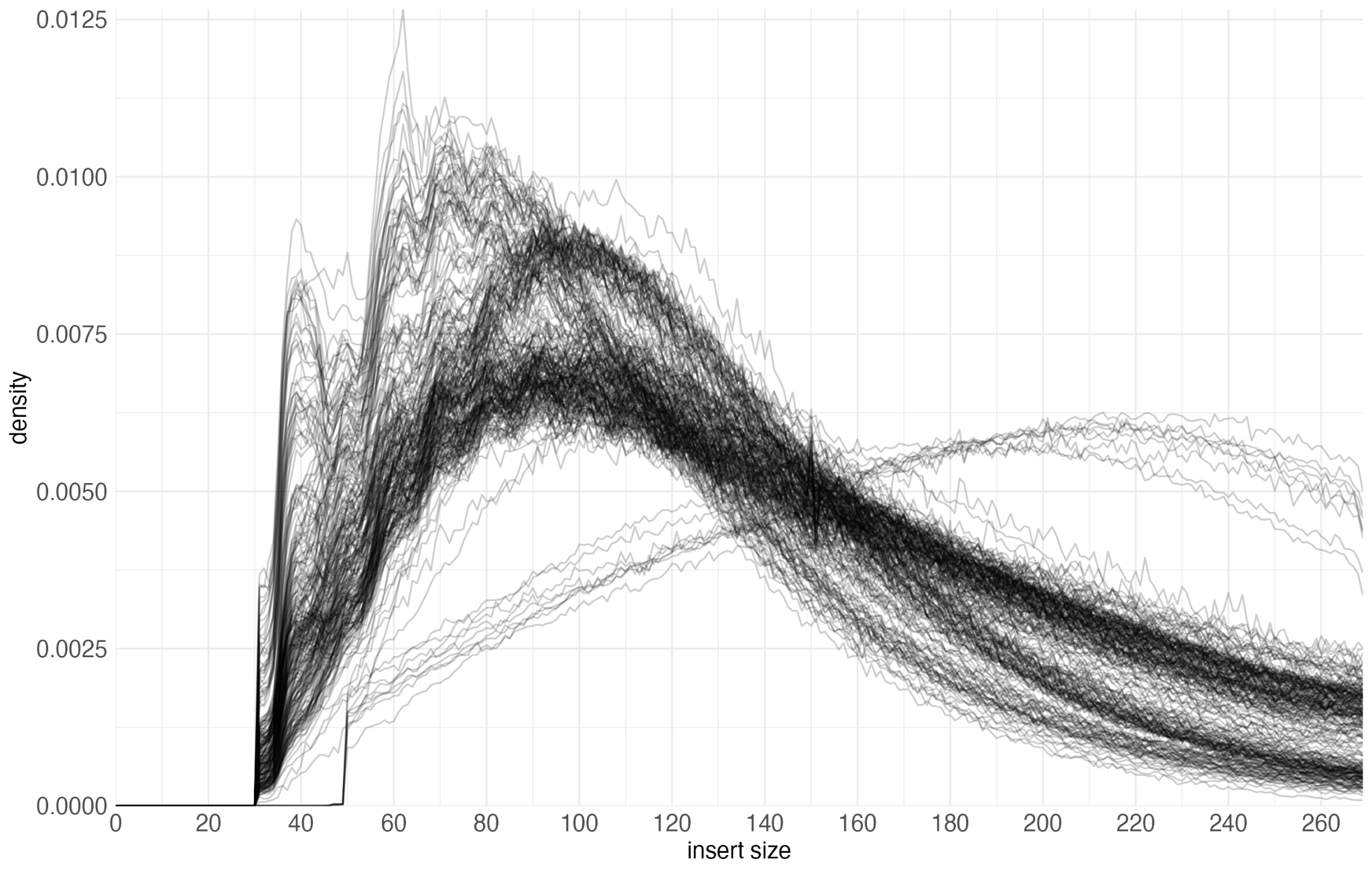

**Supplementary Figure 5.** Read length distributions for soil metagenomes from outlier metagenomes (**Supplementary Note 2**).

Hugerth, Luisa W., John Larsson, Johannes Alneberg, Markus V. Lindh, Catherine Legrand, Jarone Pinhassi, and Anders F. Andersson. 2015. “Metagenome-Assembled Genomes Uncover a Global Brackish Microbiome.” *Genome Biology* 16 (December): 279.

Pascoal, Francisco, Maria Paola Tomasino, Roberta Piredda, Grazia Marina Quero, Luís Torgo, Julie Poulain, Pierre E. Galand, et al. 2023. “Inter-Comparison of Marine Microbiome Sampling Protocols.” *ISME Communications* 3 (1): 84.

Sunagawa, Shinichi, Luis Pedro Coelho, Samuel Chaffron, Jens Roat Kultima, Karine Labadie, Guillem Salazar, Bardya Djahanschiri, et al. 2015. “Ocean Plankton. Structure and Function of the Global Ocean Microbiome.” *Science* 348 (6237): 1261359.
